# Reconstitution of *C9orf72* (GGGGCC) repeat-associated non-AUG translation with purified components

**DOI:** 10.1101/2023.05.29.542719

**Authors:** Hayato Ito, Kodai Machida, Morio Ueyama, Yoshitaka Nagai, Hiroaki Imataka, Hideki Taguchi

**Affiliations:** School of Life Science and Technology, Tokyo Institute of Technology, Yokohama 226-8503, Japan; Cell Biology Center, Institute of Innovative Research, Tokyo Institute of Technology, Yokohama 226-8503, Japan; Department of Neurology, Kindai University Faculty of Medicine, Osaka-sayama 589-8511, Japan; Graduate School of Engineering, University of Hyogo, Himeji, Hyogo 671-2280, Japan

**Keywords:** RAN translation, reconstituted cell-free translation, C9orf72, amyotrophic lateral sclerosis, ALS, GGGGCC repeat

## Abstract

Nucleotide repeat expansion of GGGGCC (G_4_C_2_) in the non-coding region of *C9orf72* is the most common genetic cause underlying amyotrophic lateral sclerosis (ALS) and frontotemporal dementia (FTD). Transcripts harboring this repeat expansion undergo the translation of dipeptide repeats via a non-canonical process known as repeat-associated non-AUG (RAN) translation. In order to ascertain the essential components required for RAN translation, we successfully recapitulated G_4_C_2_-RAN translation using an in vitro reconstituted translation system comprising human factors, namely the human PURE system. Our findings conclusively demonstrate that the presence of fundamental translation factors is sufficient to mediate the elongation from the G_4_C_2_ repeat. Additionally, we observed ribosomal frameshifting from the poly Gly-Ala dipeptide frame to other frames during the elongation process. Furthermore, the initiation mechanism proceeded in a 5’ cap-dependent manner, independent of eIF2A or eIF2D. In contrast to cell lysate-mediated RAN translation, where longer G_4_C_2_ repeats enhanced translation, we discovered that the expansion of the G_4_C_2_ repeats inhibited translation elongation using the human PURE system. These results suggest that the repeat RNA itself functions as a repressor of RAN translation. Taken together, our utilization of a reconstituted RAN translation system employing minimal factors represents a distinctive and potent approach for elucidating the intricacies underlying RAN translation mechanism.

**Significance statement:** Every cellular protein undergoes synthesis through a process known as translation. While the fundamental aspects of translation have been established, recent advancements have unveiled various noncanonical translation pathways, including the translation originating from “noncoding” RNAs. Within this context, certain neurodegenerative diseases, such as amyotrophic lateral sclerosis (ALS), are linked to the translation of noncoding RNAs, referred to as repeat-associated non-AUG (RAN) translation, the underlying mechanism of which remains controversial. To dissect the complicated nature of RAN translation, this study employs a reconstituted cell-free translation system comprised of human translation factors. By reconstituting RAN translation utilizing a minimal set of factors, this bottom-up approach not only facilitates the elucidation of its mechanism but also offers a distinctive avenue for pharmaceutical development.

## Introduction

The abnormal expansion of specific nucleotide repeats (repeat expansion) within the genome has been established as a causative factor in neurodegenerative diseases such as amyotrophic lateral sclerosis (ALS) and spinocerebellar ataxia (SCA) (1, 2). Recent investigations have revealed a noncanonical translation mechanism for transcripts containing repeat expansions, termed repeat-associated non-AUG (RAN) translation, which operates independently of the conventional initiation codon AUG (3–9). Notably, a representative instance of RAN translation occurs at the GGGGCC (G_4_C_2_) repeat located in the first intron of *C9orf72* (Fig. S1A) (referred to as C9-RAN), constituting the most prevalent genetic mutation observed in familial ALS (10, 11). C9-RAN involves the translation of all conceivable frames of the G_4_C_2_ repeat, which is transcribed in both directions. Specifically, the sense strand (GGGGCC) yields dipeptide repeats (DPRs) consisting of Gly-Ala (GA from GGG-GGC, 0 frame), Gly-Pro (GP from GGG-CCG, +1 frame), and Gly-Arg (GR from GGC-CGG, +2 frame). Indeed, these DPRs have been detected not only in patient tissues (4, 6, 12, 13), but also in diverse model organisms, and their association with cytotoxicity is well-documented (4, 6, 13–20). Based on these recent findings, it is proposed that C9-RAN may assume a central role in the pathogenesis of repeat expansion-associated ALS, offering a novel avenue to explore potential therapeutic targets for the disease.

The translation process consists of four fundamental steps: initiation, elongation, termination, and ribosome recycling (21). In eukaryotes, the canonical initiation of translation takes place through a scanning mechanism mediated by the cap structure at the 5’-end of the mRNA (21–23). This scanning mechanism is strictly regulated by over ten translation initiation factors in conjunction with the eIF4F complex, comprising eIF4A, eIF4G, and eIF4E, which initiates translation from the canonical AUG initiation codon. Recent genome-wide studies have unveiled diverse instances of non-AUG translation, including those initiated from CUG and GUG codons (24–26). Non-AUG translation is believed to involve eIF2A and eIF2D, alternative factors for eIF2 (27–30). Furthermore, it is recognized that certain eukaryotic mRNA and some genes within viral genomes commence translation without scanning, utilizing a sophisticated RNA structure element termed an internal ribosomal entry site (IRES) (23, 31).

The initiation mechanism of C9-RAN exhibits commonalities with canonical mechanisms. For example, previous studies utilizing monocistronic reporters have shown that C9-RAN initiates from near cognate codons, such as CUG and AGG, located upstream of the G_4_C_2_ repeat sequence, through a scanning mechanism involving eIF4A (32–34). However, contrasting findings employing bicistronic reporters have indicated that C9-RAN commences independent of the cap structure (35, 36). Furthermore, diverse mechanisms generating multiple translational frames have been reported, such as ribosomal frameshift during elongation on the repeat sequence and initiation from distinct start codons for each frame (33, 34, 37, 38). The occurrence of frameshift in C9-RAN is predicted to be associated with a secondary structure of the mRNA, namely the guanine quadruplex (G4-RNA), formed as a consequence of the guanine-rich property of the G_4_C_2_ repeat (33). In addition, it has been observed that C9-RAN is regulated by factors that are not essential for canonical translation. Notably, eIF2A, an alternative factor for eIF2, has been postulated to modulate C9-RAN, although its requirement in transfected cells remains a topic of controversy (35, 39, 40).

Hence, numerous discussions ensue regarding the molecular mechanism of C9-RAN, primarily attributed to the lack of consistent findings concerning the fundamental process of translation and factors associated with C9-RAN. The utilization of cell-based experimental systems, such as transfected cells and cell lysate translation systems, is believed to contribute to these inconsistencies. Such cell-based systems encompass proteins unrelated to translation and exhibit varying quantities of translation factors contingent upon cell types (41), thereby yielding a heterogeneous array of results, occasionally conflicting in nature. Indeed, GP frames occurring in C9-RAN have been shown to exhibit different translation efficiencies depending on the cell type (32, 33). Consequently, to ascertain the intricate molecular mechanism of C9-RAN comprehensively, the employment of an in vitro translation system independent of cell lysate becomes imperative.

To circumvent such complexities associated with C9-RAN and elucidate its fundamental, the adoption of a reductionist approach is invaluable. In this context, a cell-free translation system reconstituted in vitro, employing purified factors indispensable for translation, serves as the optimal experimental system for conducting a comprehensive investigation of the mechanism governing C9-RAN. Subsequent to the establishment of a reconstituted translation system in *E. coli* (42), recent advancements have led to the development of eukaryotic translation factor-based reconstituted translation systems (43–45). Within the human factor-based reconstituted translation system (human PURE, Fig. S2), a reconstituted system using IRES (human PURE-IRES) (43), thereby omitting initiation factors, was first developed. This was succeeded by the development of a fully reconstituted system, enabling translation initiation in a cap-dependent manner in the presence of numerous initiation factors (human PURE-cap) (44).

In this study, we successfully recapitulated C9-RAN using the human PURE system. By employing distinct modes of translation, namely IRES-dependent and cap-dependent, we were able to distinguish and investigate the elementary processes underlying C9-RAN. Through the utilization of this minimalist system, we acquired valuable insights into translation elongation, ribosomal frameshifting, and initiation within the context of C9-RAN. Furthermore, we observed the suppression of C9-RAN with longer repeat RNA in human PURE. These findings shed light on previously ambiguous aspects of the molecular mechanism underlying C9-RAN, highlighting the merits of employing the bottom-up approach facilitated by human PURE as a versatile tool for dissecting RAN translation associated with other diseases.

## Results

### G4C2 translation elongation with minimal translation factors

To investigate the elementary steps involved in C9-RAN, we employed human PURE system, including purified ribosomes, eukaryotic initiation/elongation/release factors (eIFs, eEFs, and eRFs) (43, 44) (Fig. 1A, Fig. S2). Initially, we aimed to determine whether the G_4_C_2_ repeat sequences could undergo elongation with the minimal translation elongation factors. For this purpose, we used the hepatitis C virus (HCV) IRES to recruit the ribosome in the absence of eIFs (46). We introduced the HCV IRES, along with the ATG codon following the IRES, upstream of the GA frame (GGG-GCC/ 0 frame) within the G_4_C_2_ 80 repeat (80R) sequence (Fig. 1B) as previous studies have indicated that the translation product derived from the GA frame is the most abundant in C9-RAN (6, 32, 33). We conducted translation experiments using a fusion gene encoding HCV-IRES-ATG-G_4_C_2_-80R, followed by a Myc-tag for detection, in HeLa lysate and the human PURE-IRES. We detected translation products of the expected molecular weight (∼ 25 kD for ATG-G_4_C_2_ 80R-Myc) in both HeLa lysate and the human PURE-IRES, only when IRES was present (Fig. 1C), although the translation product yield was considerably smaller in the human PURE-IRES. Given that this experiment forced the translation initiation to the IRES-linked ATG codon, this translation does not meet the definition of RAN translation. Nevertheless, this experiment is important in that it extracted only the elongation step in translation and demonstrates that the G_4_C_2_ repeat can undergo elongation with minimal elongation factors.

**Fig. 1:**
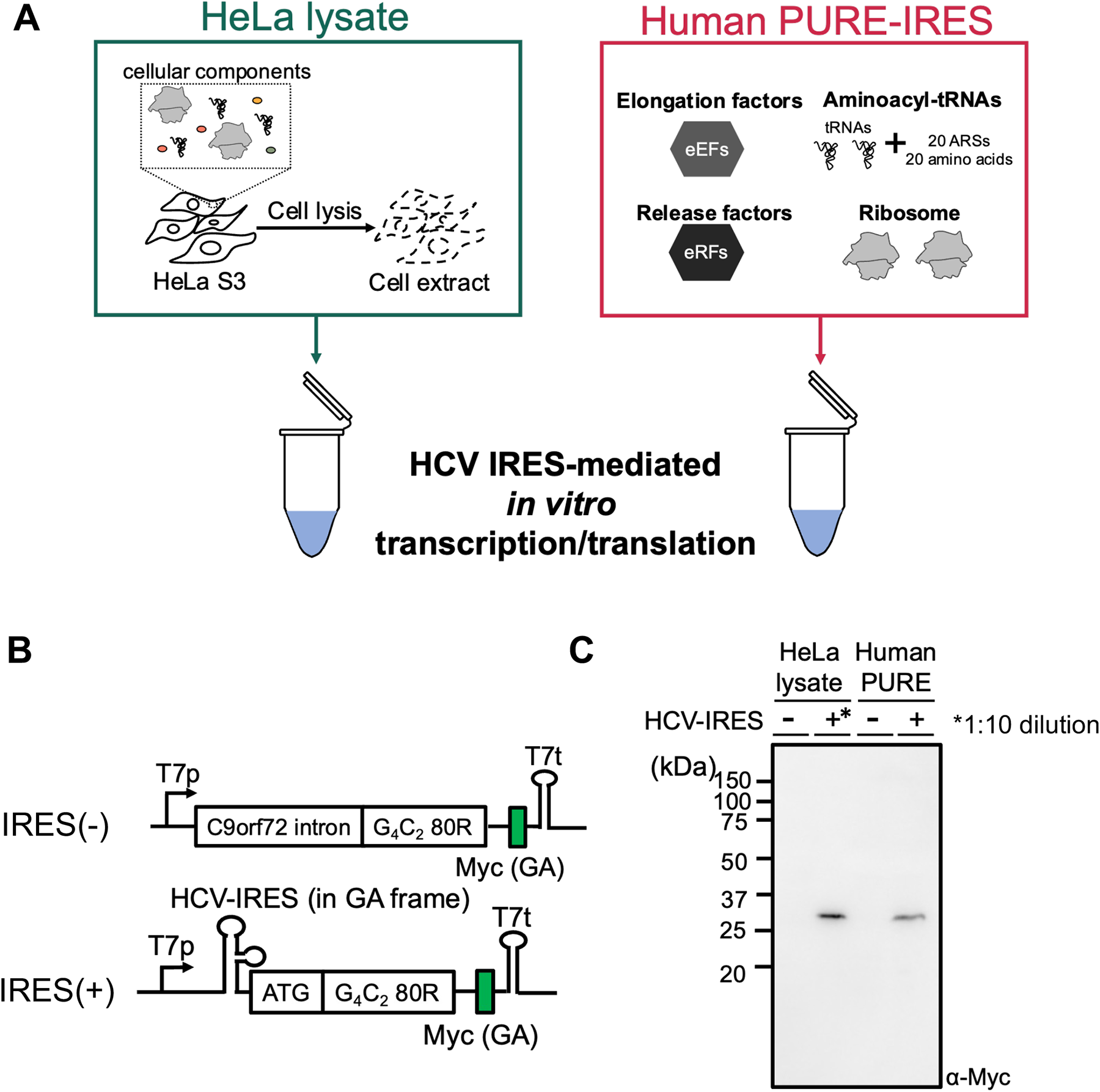
Minimal translation factors are sufficient for translation of the G_4_C_2_ repeats. (*A*) Illustration of each in vitro translation system. HeLa lysate (63) contained intracellular factors, while the human PURE-IRES system utilized purified translation factors (43). Hepatitis C Virus (HCV)-IRES, enabling translation initiation without initiation factors (46), was employed to investigate the elongation mechanism of the G_4_C_2_ repeats. (*B*) Schematic representation of the construct for monitoring the elongation of the G_4_C_2_ repeats. HCV-IRES was inserted upstream of the G_4_C_2_ repeat sequence, with a Myc-tag introduced downstream of the G_4_C_2_ repeat in the GA frame. (*C*) Anti-Myc western blot of the G_4_C_2_ repeat reporter plasmids expressed in each in vitro translation system. To prevent band detection saturation in HeLa lysate, IRES(+) reaction was diluted 1:10 in the sample buffer, as indicated by the asterisk (*).

### Ribosomal frameshifting during elongation in the GA (0) Frame

Next, we established a nano-luciferase (Nluc) reporter assay system to quantitatively measure the translation products with high sensitivity. Nluc and an HA-tag were introduced downstream of the IRES-ATG-(G_4_C_2_)_80_ or IRES-TTT-(G_4_C_2_)_80_ genes (Fig. 2A). In addition to the GA (0) frame, we generated constructs with two additional reading frames: the GP (+1) frame and the GR (+2) frame, achieved by inserting one or two nucleotides, respectively, following the (G_4_C_2_)_80_ repeat. Western blotting analysis following the translation of the Nluc constructs using the human PURE-IRES or HeLa lysate demonstrated an ATG-dependent expression of the GA frame product compared to the products from other frames (Fig. 2B, D). The ATG-dependency, along with the approximate expected molecular weight for the ATG-(G_4_C_2_)_80_-Nluc-HA gene (∼39 kD), indicates that translation elongation commences from the ATG immediately after the IRES. Notably, translation of the TTT-containing gene using the HeLa lysate yielded a small amount of the 39 kD protein in the GA frame (Fig. 2B), underscoring the utilization of the TTT codon as a noncanonical start codon.

**Fig. 2:**
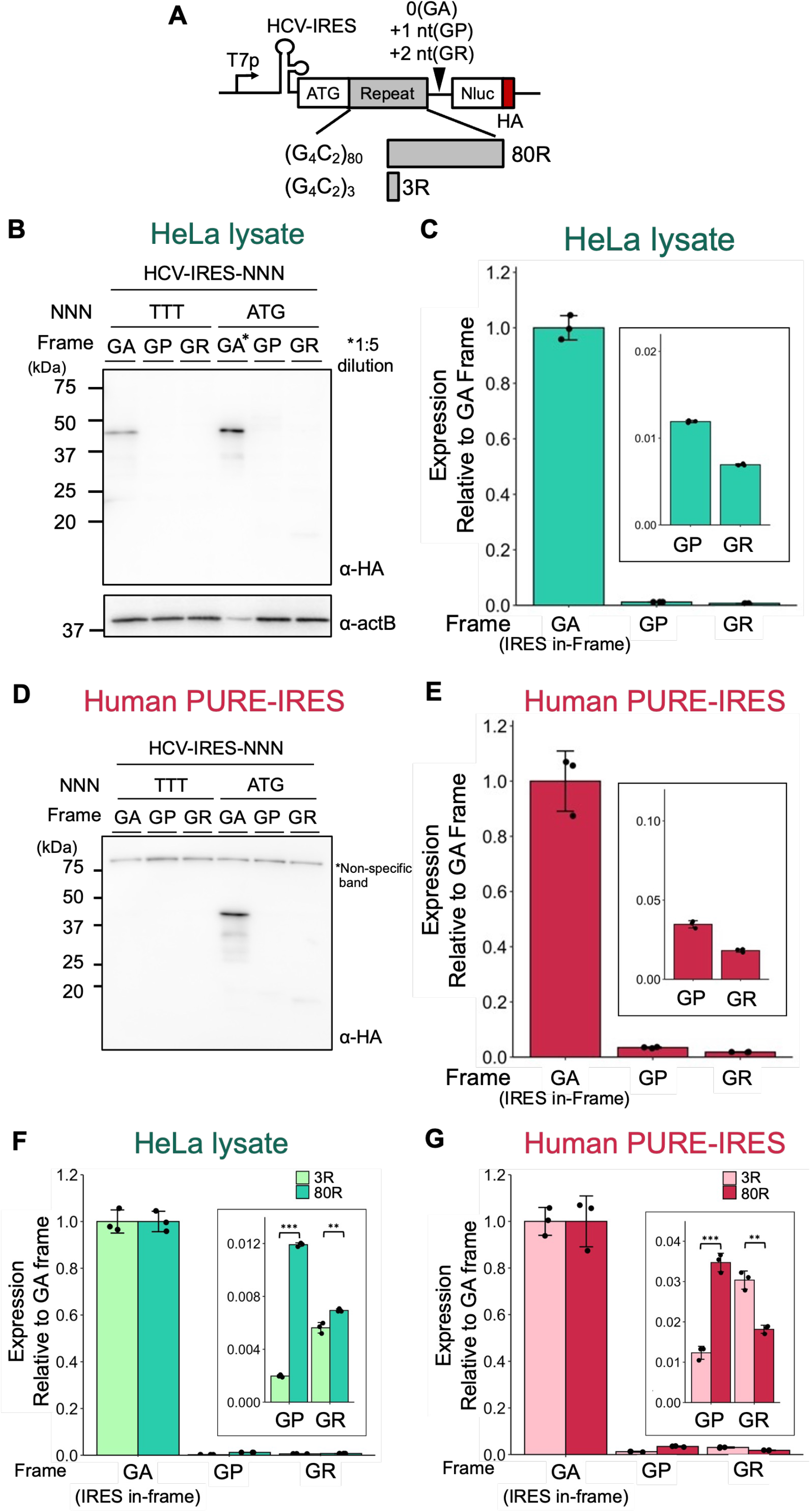
Ribosomal frameshift occurs in the GA (0) frame. (A) Schematic representation of the elongation reporters for the G_4_C_2_ repeats. Nano-Luciferase (Nluc) was used as a reporter enzyme to quantify translation efficiency. HA-tag in the GA-frame was introduced downstream of the G_4_C_2_ repeat. (*B, D*) Anti-HA western blot of the elongation-reporter plasmids expressed in each in vitro translation system. (*B*) HeLa lysate, (*D*) human PURE-IRES. (*C, E*) Relative expression from IRES-ATG-(G_4_C_2_)_80_-Nluc-HA normalized to the GA frame. (*C*) HeLa lysate, (*E*) human PURE-IRES. (*F-G*) Expression of IRES-ATG-(G_4_C_2_)_3_-Nluc and IRES-ATG-(G_4_C_2_)_80_-Nluc normalized to their respective GA frames. (F) HeLa lysate, (G) human PURE-IRES. Error bars represent standard deviations (±SD) from three technical replicates. Two-tailed student’s *t* test, \*\**p*≦0.01, \*\*\**p*≦0.001. Raw data of the Nluc assay are provided in Supplementary Table S1.

We first confirmed that luciferase activity-based translation was entirely dependent on the IRES in both HeLa lysate and the human PURE-IRES translation (Fig. S3A). Subsequently, we reproducibly observed luciferase activities across all frames, including the minor GP (+1) and GR (+2) frames, during the translation of the genes containing either ATG or TTT start codons, using both HeLa lysate and the human PURE-IRES (Fig. 2C, E, Fig. S3B, C). Earlier studies employing lysate-based translation also showed translation occurring in all frames (32, 33), with the GP and GR frames suggested to result from ribosomal frameshifting originating from the GA frame (33). Through a series of experiments using the human PURE-IRES, our observation of translation initiation from the ATG or TTT start codons immediately after the IRES (Fig. 2D, E, Fig. S3C), along with the minimal translation activity in the absence of IRES (Fig. S3A), led us to suggest the occurrence of frameshifting even within a simplified translation system. The luciferase activities of the GP and GR frames relative to the GA frame in ATG-mediated human PURE-IRES translation were 3.5% and 1.9%, respectively (Fig. 2E), higher than those observed in HeLa translation (1.2% and 0.7% for GP and GR, respectively, Fig. 2C). This suggests the possible presence of a frameshift repressor in the lysate.

Then, we examined the potential influence of the G_4_C_2_ repeat length on the translation of GP (+1) and GR (+2) frames in the human PURE-IRES, considering that a prior study using lysate translation demonstrated that frameshifting on the G_4_C_2_ repeat sequence is enhanced by longer repeat RNA (33). Within the human PURE-IRES, translation of the GP frame exhibited an increase, while the GR frame displayed a decrease in response to longer repeat lengths (Fig. 2G), thus indicating that the length of the repeat exerts an influence on frameshifting.

### Translation initiation in C9-RAN can be initiated using minimal initiation factors

Subsequently, we investigated the initiation mechanism of C9-RAN by employing a fully reconstituted 5’ cap-dependent translation system equipped with canonical thirteen eIFs (referred to as human PURE-cap, Fig. 3A) (44). In order to quantify C9-RAN via a scanning mechanism, we constructed a series of reporter mRNAs, utilizing the methodology previously developed by Green *et al* (32). Initially, we evaluated the property of human PURE-cap in non-AUG translation, employing mRNAs containing CUG or AGG codons fused with Nluc. We observed a relatively higher translation efficiency for the CUG codon compared to that observed in HeLa lysate (Fig. S4). Then, we proceeded to translate a reporter mRNA harboring a 5’-cap, the *C9orf72* intron, (G_4_C_2_)_80_ repeats, followed by Nluc and a FLAG-tag (Fig. 3B). Through Western blotting analysis, we confirmed the synthesis of ∼40 kD protein bands corresponding to the GA frame in both the human PURE-cap system and HeLa lysate (Fig. 3C, Fig. S5A). The approximate molecular weight of these protein bands aligns with the expected size (35 kD) if translation was initiated from the upstream sequence of the repeats, as previously demonstrated (32–35). This result, obtained using the human PURE-cap system, provides evidence that C9-RAN can be initiated utilizing minimal translation factors. Additionally, translation in HeLa lysate produced a ∼23kD protein band in the GR frame (Fig. S5A). The molecular weight of ∼23 kD corresponds well with the calculated size (23.5 kD) of the translation product when initiation occurs at the downstream AUG codon within the G_4_C_2_ repeats in the GR frame, indicating that this product likely represents a leaky scanning product (Fig. S5B).

**Fig. 3:**
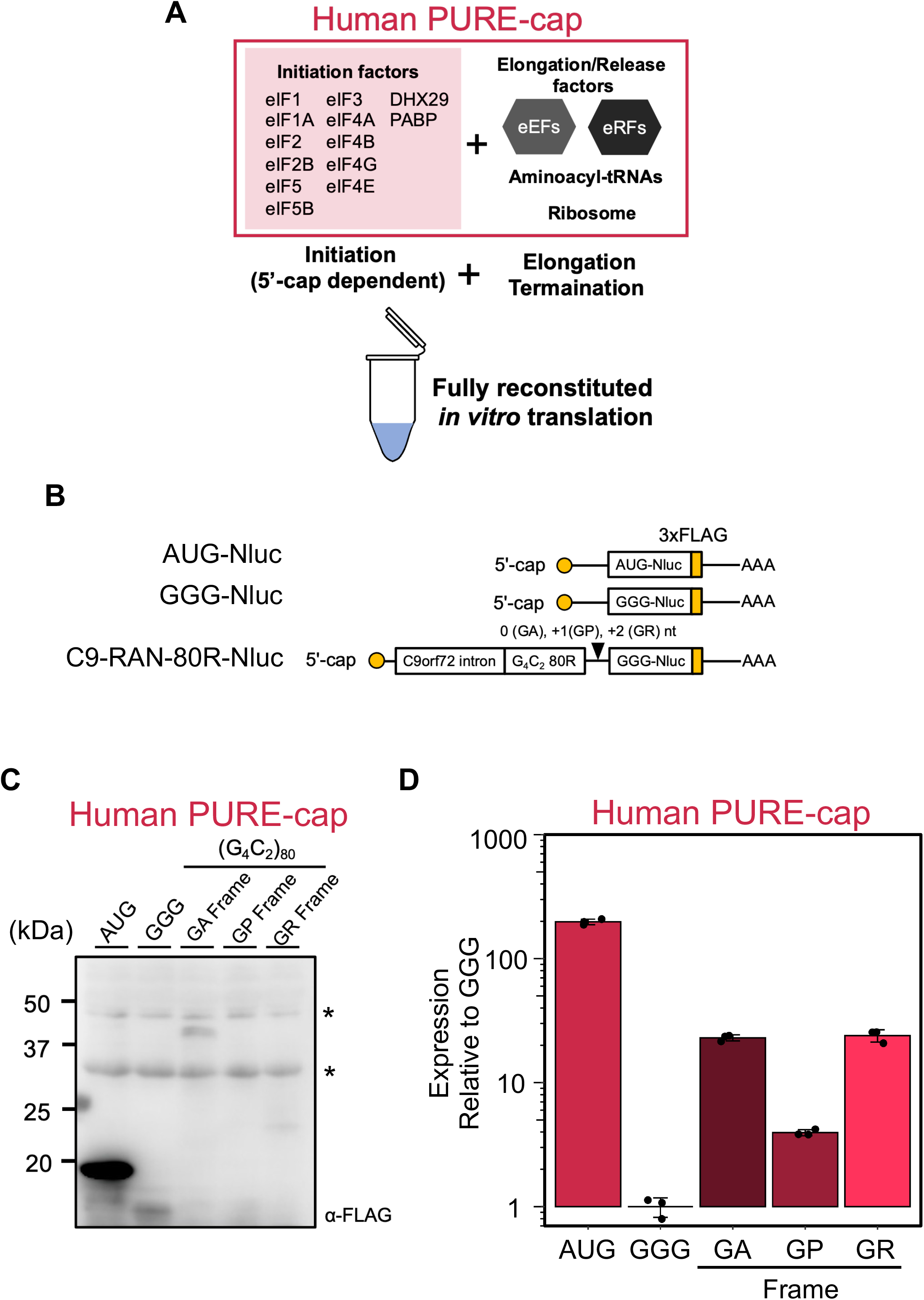
Minimal translation factors are enough for initiating C9-RAN translation. *(A)* Illustration depicting the human PURE-cap system (44). *(B)* Diagram presenting the C9-RAN reporters based on previously reported designs (32). Nluc was used employed the reporter enzyme to quantify translation efficiency. A FLAG-tag was inserted downstream of the G_4_C_2_ repeat in each frame. *(C)* Anti-FLAG western blot of the C9-RAN reporter plasmids expressed in human PURE-cap. An asterisk (*) indicates a non-specific band. *(D)* Relative expression of the C9-RAN reporters normalized to GGG-Nluc in human PURE-cap. Error bars represent ±SD from three technical replicates. Raw data of the Nluc assay are provided in Supplementary Table S2.

Luciferase activities were detected in all frames during both HeLa lysate and human PURE-cap translation, strictly dependent on the presence of a cap structure (Fig. 3D, Fig. S5C. Fig. S6A), thereby confirming the sufficiency of minimal translation factors for C9-RAN. However, the overall translation level of C9-RAN in the human PURE was lower compared to that in HeLa lysate. For instance, the translation of the GA frame in HeLa lysate was comparable to that of AUG-Nluc mRNA (Fig. S5C), whereas it constituted approximately one-tenth of the translation observed in human PURE-cap (Fig. 3D). This discrepancy suggests that cap-dependent translation using the PURE system requires additional factors to enhance the efficiency of translation. Moreover, the deletion of the *C9orf72* intron significantly suppressed translation initiation of the GA frame in both human PURE and HeLa lysate (Fig. S6B), indicating that the *C9orf72* intron serves as the primary initiation region in the GA frame even during translation mediated by the human PURE.

We subsequently investigated the putative start codon in human-PURE translation. Considering that the CUG codon within the *C9orf72* intron has been proposed as the start codon in the GA frame (Fig. S1) (32–35), we introduced mutations to replace CUG with AUG or GGG (Fig. S6C). The AUG substitution significantly enhanced the expression of the GA frame in both HeLa lysate and Human PURE-cap, whereas the GGG codon markedly decreased the expression of the GA frame in both HeLa lysate and human PURE-cap (Fig. S6C). These findings, obtained through the use of human PURE, provide compelling evidence that C9-RAN initiation occurs from the CUG codon within the *C9orf72* intron in a cap-dependent manner, independent of any *trans*-acting factors beyond essential translation factors.

### C9-RAN translation is suppressed by longer G4C2 repeats in human PURE translation

Enhancement of RAN translation can be attributed to longer repeat RNA sequences (32–35, 47–49). Given the lack of comprehensive understanding regarding the underlying mechanism, we investigated the dependence of repeat-length in human PURE translation (Fig. 4A). In the HeLa lysate translation, the augmentation of C9-RAN occurred in accordance with the length of G_4_C_2_ repeats in both GA and GP frames (Fig. 4A, Fig. S7). This enhancement reached saturation at 29 repeats, exhibiting an approximately nine-fold increase in comparison to the three-repeat scenario in the GA frame (Fig. 4A). Conversely, in contrast to the HeLa lysate translation, the utilization of human PURE-cap translation inhibited C9-RAN from 29 repeats, leading to decline of less than 10% in the GA frame translation when utilizing 80 repeats (Fig. 4A). The inhibition of repeat length was also observed in the GP frame during human PURE-cap translation (Fig. S7).

**Fig. 4:**
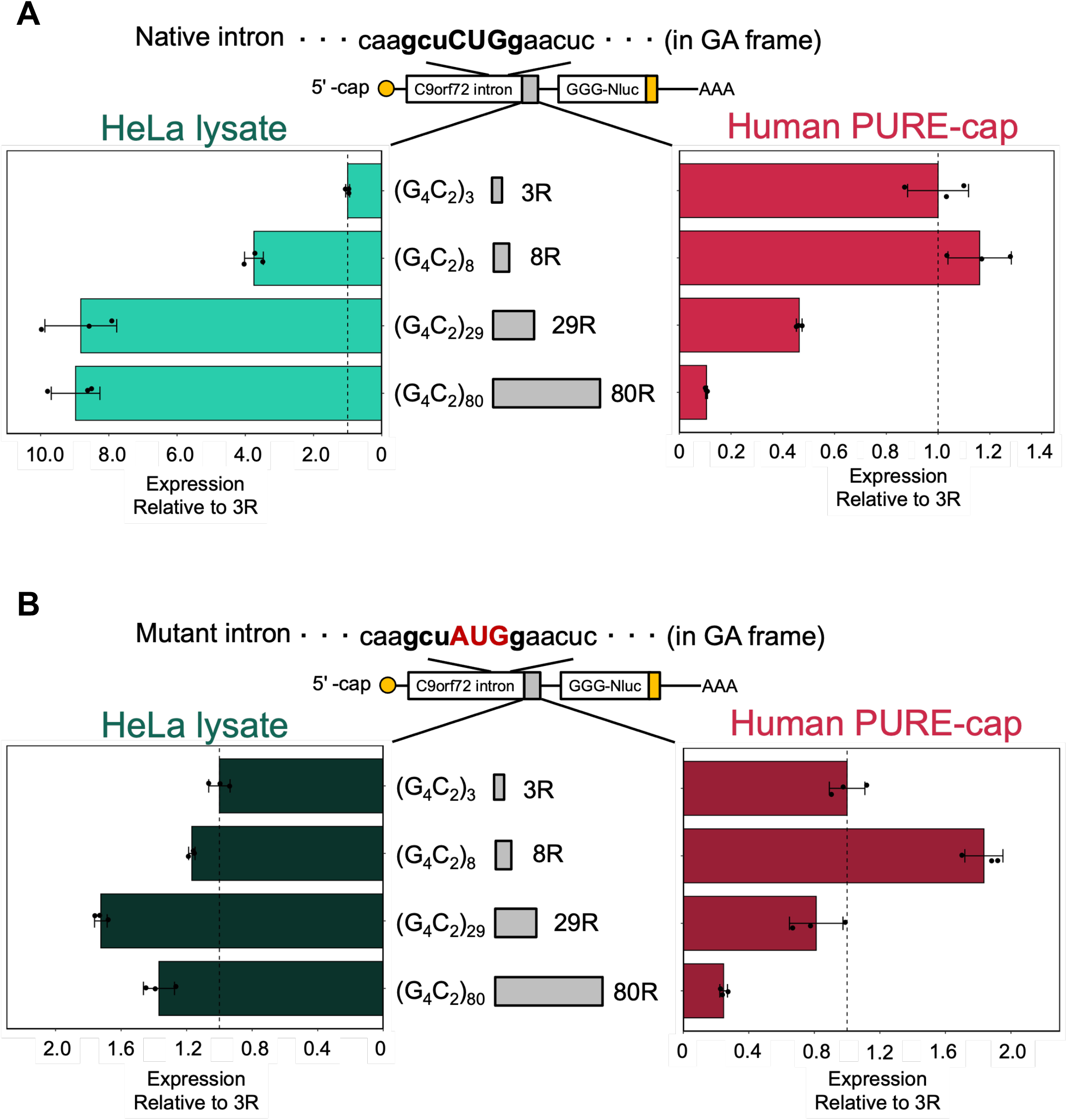
Inhibition of C9-RAN by longer repeat RNA in human PURE-cap. (*A, B*) Relative expression levels of C9-RAN reporters with varying numbers of repeats in each in vitro translation system. Normalized to 3R-Nluc. (*A*) The first intron of *C9orf72*. (*B*) Mutant intron substituting CUG with AUG codon. The CUG codon located -24 nt upstream of the repeat sequence is predicted as the GA frame start codon (32–35). Error bars represent ±SD from three technical replicates.

We hypothesized that the length-dependent inhibition observed in the reconstituted translation system could be attributed to aberrant non-AUG initiation in human PURE-cap, as essential eukaryotic initiation factors (eIFs) such as eIF2A and eIF2D, known to be involved in non-AUG translation initiation (28–30, 50), are absent. To test this hypothesis, we introduced a CUG-to-AUG substitution in the *C9orf72* intron. Interestingly, this substitution nearly abolished the length-dependent enhancement observed in the HeLa lysate translation (Fig. 4B). However, in human PURE-cap, the CUG-to-AUG substitution did not alter the declining trend in translation efficiency observed beyond 29 repeats (Fig. 4B), suggesting that the longer repeat RNA-mediated suppression of C9-RAN in human PURE-cap is not solely attributed to an anomaly in non-AUG initiation.

## Discussion

In this study, we successfully RAN translation from *C9orf72*-(G_4_C_2_) repeat (C9-RAN) using a reconstituted translation system comprised of purified human translation-related factors (human PURE). The pivotal and straightforward message conveyed by this study is the explicit demonstration that C9-RAN occurs exclusively through the utilization of canonical translation factors. We particularly emphasize the unique advantage of human PURE in dissecting elementary steps of translation, including elongation and initiation.

Before discussing the details of the specific findings regarding these elementary steps, it is important to acknowledge certain limitations and considerations associated with the use of human PURE. Firstly, while it has been established that C9-RAN can proceed with minimal factors, it does not imply that such minimal factors are representative of physiological conditions. The results presented here do not rule out the possibility that the presence of additional factors might alter the properties of C9-RAN, either qualitatively or quantitatively. Indeed, it is worth noting true that translation efficiency in the human PURE was lower compared to that in the HeLa lysate system. Hence, it is plausible that the accumulation of translation products leading to toxicity may necessitate the involvement of supplementary factor(s).

Through the use of human PURE-IRES, which skips the complicated initiation reaction, we unequivocally demonstrated that the G_4_C_2_ repeat is translated by a defined set of eEFs. Furthermore, by enforcing translation initiation exclusively from the ATG codon immediately following the IRES, we were able to evaluate frameshifting from the GA (0) frame, as discussed in further detail below.

There have been debates regarding the requirement of a 5’ cap structure for C9-RAN (32, 33, 35, 36). Human PURE-cap experiments definitively established that the translation initiation mechanism for C9-RAN is indeed 5’ cap-dependent. These results not only corroborate previous studies that demonstrated 5’ cap-dependent C9-RAN using reporter assays, including those employing cell lysates (32, 33), but also suggest that even within a cellular environment where numerous factors are present, a specific set of eIFs could play a primary role in C9-RAN.

C9-RAN initiation, potentially involving canonical eIFs like eIF2, has been proposed (2, 32, 33, 39). Considering the specific nature of non-AUG translation in C9-RAN, it is plausible that eIF2A and eIF2D play roles in translation initiation. Indeed, previous studies implicated eIF2A and eIF2D as regulators of C9-RAN (35, 39, 40), and another regulator of non-AUG translation, 5MP, has also been linked to RAN translation (51). It is noteworthy that the findings from human PURE-cap experiments indicate that eIF2A and eIF2D are not involved in the initiation of C9-RAN.

An intriguing and unresolved aspects of RAN translation, including C9-RAN, is the generation of translation products originating from multiple frames (2, 52, 53). Previous studies conducted in cultured cells and cell lysates could not distinguish whether the translation of multiple frames results from ribosomal frameshifting or the presence of multiple translation initiation sites (33). Through the dissection of C9-RAN using human PURE, we revealed that both mechanisms contribute to this phenomenon. Firstly, analysis utilizing human PURE-IRES uncovered the occurrence of frameshifting in the GA frame (Fig. 2). Furthermore, this analysis demonstrated that longer G_4_C_2_ repeats enhance frameshifting in the GP frame. We propose that this enhancement, mediated by longer repeats, is attributed to the formation of G4RNA structures within the G4C2 repeats, as previous research indicates that G4RNA suppresses ribosomal translocation and promotes frameshifting (54). It is worth noting that the frameshift efficiency in HeLa lysate was lower than that observed in human PURE-IRES. Therefore, the presence of specific factors in HeLa lysate may suppress ribosomal frameshifting. For instance, the abundance of RNA-binding proteins, such as TDP-43, known as RNA chaperones (55), might lead to alterations in the G4RNA structure. Additionally, the interaction of the ribosome with Shiftless, a factor that suppresses -1 programmed frameshift (56), may also influence the frameshift process.

Utilizing the human PURE-IRES system, we observed a ribosomal frameshift from the GA frame to the GP frame at a frequency of 3.5% (Fig. 2). In contrast, 5’ cap-dependent translation initiation using human PURE-cap exhibited higher efficiency in generating the GP frame, with a ratio of +1 to 0 frame of approximately 17% (Fig. 3). This outcome suggests the involvement of an alternative mechanism that leads to the initiation at different sites, giving rise to the production of multiple frames, including the GP frame. Considering that human PURE-IRES and human PURE-cap differ solely in the presence of eIFs, approximately 80% (3.5%/17%) of the +1 frame may undergo scanning-specific translation initiation mediated by the 5’ cap and canonical eIFs. Nevertheless, the initiation site for GP frame translation remains elusive. As the GP frame contains a stop codon preceding the G_4_C_2_ repeat (Fig. S1), it is plausible that initiation occurs from within the repeat sequence (Fig. S6B). Furthermore, G4RNA has been demonstrated to induce read-through of the stop codon (54), implying that multiple translation mechanisms could contribute to GP frame translation.

Our analysis using human PURE-cap revealed a striking difference in repeat-length dependency. Cap-dependent C9-RAN was inhibited by longer repeat RNA in human PURE-cap. While previous reports have indicated that C9-RAN is enhanced in a repeat length-dependent manner in cell lysate translation systems and transfected cells (33, 48), our findings suggest the involvement of non-canonical translation factors in the repeat length-dependent stimulation of C9-RAN. In the human PURE-cap system, translation increase was observed for the GA frame up to (GGGGCC)_8_ repeats; however, translation efficiency was significantly reduced beyond 29 repeats, which are likely to form the G4RNA structure. Additionally, the mutation of the initiation codon CUG to the AUG codon in the GA frame also displayed inhibitory tendencies. Therefore, it is plausible that the inhibition of longer repeats-induced C9-RAN arises from elongation inhibition, potentially caused by the formation of G4RNA. Indeed, previous studies have demonstrated that non-AUG translation, including RAN translation, can be facilitated by downstream secondary structures (57, 58).

If the characteristics of the repeat RNA itself contribute to the length-dependent inhibition, it is necessary to consider factors that bind to the RNA repeats. Alongside the RNA chaperone, which modulates RAN translation by interacting with repeat RNA and modifying its structure (55), RNA helicases that unwind the higher-order structure of repeat RNA (59–61) are also likely to be involved in the process. In HeLa lysate, RNA helicases such as DHX36 are responsible for unwinding G4RNA. DHX36 has been identified as a binding partner of G4RNA and is capable of unwinding its structure in an ATP-dependent manner, thereby rescuing ribosomal stalling on the G_4_C_2_ repeat (59, 60). However, knockdown or knockout of DHX36 only results in approximately 50% inhibition, even at G_4_C_2_ 70 repeats (59), which falls significantly short of the inhibitory effect observed in human PURE-cap. Hence, it is reasonable to speculate that additional factors beyond DHX36 are involved in the elongation process of RAN translation.

The dissection analysis conducted using human PURE, in conjunction with previous studies, suggests that translation of the G_4_C_2_ repeats represents a delicate balance between the promotion of noncanonical initiation and the inhibition of repeat elongation. Consequently, the presence of a novel regulator of RAN translation that enhances ribosomal translocation and exhibits ribosomal helicase activity may be implicated in C9-RAN within the cellular context.

Lastly, the human PURE system not only serves as an invaluable tool for studying C9-RAN, but also for investigating RAN translation involving other nucleotide repeats and, more broadly, for facilitating bottom-up approaches to noncanonical translation. Furthermore, the elucidation of the underlying mechanisms governing the fundamental process of RAN translation, as revealed by this reconstituted system, holds significant potential in the development of therapeutic strategies targeting neurodegenerative diseases associated with RAN translation.

## Materials and Methods

### Plasmids

To generate the HCV-IRES-(G_4_C_2_)_80_-Myc reporter plasmid, we employed the following procedure. The HCV-IRES region was amplified from the HCV-IRES sequence and introduced upstream of the previously published pcDNA^TM^5/FRT-T7-*C9orf72* intron1-(G_4_C_2_)_80_-Myc vector (62) by means of HindIII and BssHII. In order to construct the *C9orf72* intron1-(G_4_C_2_)_80_-Nluc-3xFLAG, the Nluc-3xFLAG region was amplified from the Nluc sequence and inserted downstream of the aforementioned pcDNA^TM^5/FRT-T7-*C9orf72* intron1-(G_4_C_2_)_80_ vector using PstI. A similar strategy was employed to engineer the C9-RAN Nluc reporter with HCV-IRES. Repeat sequences of varying lengths were randomly obtained through PCR. The insertion of ATG, GGG, and near-cognate codons to the Nluc plasmid was achieved using standard cloning procedures and Gibson Assembly. Supplementary Tables S3 and S4 provide a comprehensive list of the plasmids and oligonucleotides employed in this study.

### In vitro transcription

The reporter plasmids were linearized with XbaI (Takara). Subsequently, the linearized DNA was purified using the Wizard® SV Gel PCR purification kit (Promega). For the synthesis of 5’-capped mRNA, the reactions were carried out following previously established protocols (62). Uncapped RNAs were transcribed in vitro using the CUGA T7 in vitro transcription kit (Nippon Gene). The size and quality of the resulting mRNAs were assessed through denaturing RNA gel electrophoresis.

### Human PURE in vitro translation

The HCV-IRES-dependent in vitro translation was conducted according to previously described methods (43). In brief, for the luminescence assay of nano luciferase, a mixture of human PURE cocktail (4.5 µL) and reporter DNAs (0.5 µL, 15 nM) was incubated at 32 °C for 3 h, followed by termination through incubation on ice. The subsequent steps were performed using 96-well plates. A volume of 2 µL of the samples was diluted in Glo lysis buffer (Promega) and incubated in a 1:1 ratio for 3 min in the dark with NanoGlo substrate, freshly diluted a 1:50 ratio in NanoGlo buffer (Promega), with shaking. Luminescence was measured using a Varioskan LUX Multimode Microplate Reader (Thermo Fisher Scientific). For western blotting, the same procedure was followed, except that 6 µL of samples were used. The 6 µL reactions were mixed with 15 µL of 4xSDS sample buffer (240 mM Tris-HCl pH 6.8, 40% (v/v) glycerol, 0.01% (w/v) bromophenol blue, 7% (w/v) SDS, 10% (v/v) of 2-mercaptoethanol) and heated at 70 °C for 10 min. Subsequently, 20 µL of the samples were loaded onto a 13% SDS-polyacrylamide gel for electrophoresis and subsequent western blotting analysis.

The cap-dependent in vitro translation was conducted following the established protocol (44). Briefly, the human PURE-cap cocktail (3.6 µL) was combined with 0.4 µL of reporter mRNAs (final concentration, 60 nM) and incubated at 32 °C for 6 h. The samples were analyzed using the same approach as the HCV-IRES-dependent human PURE translation.

### HeLa lysate in vitro translation

The in vitro translation system derived from HeLa S3 cells was as previously described (63). The HCV-IRES-dependent translation was assessed using the same procedure as the human PURE translation after incubation at 32 °C for 2 h. In the cap-dependent in vitro translation, reporter mRNAs were added at a concentration of 3 nM and incubated at 32 °C for 1 h. The samples were analyzed following the same method as the human PURE in vitro translation.

### Western Blotting

The samples were separated on 13% SDS-polyacrylamide gels and transferred onto PVDF membranes. The membranes were blocked with 2% (w/v) skim milk in TBS-T. Subsequently, the membranes were incubated with primary antibodies listed in Supplementary Table S5. After washes with TBS-T, the membranes were incubated with HRP-conjugated anti-mouse IgG (Sigma-Aldrich). Chemiluminescence signals were detected using an LAS4000 (FujiFilm)

### Quantification and statistical analysis

All statistical analyses were performed using custom R code. The presented quantitative data represent the mean ± S.D. from a minimum of three independent experiments.

## Acknowledgments

We thank Biomaterials Analysis Division, Open Facility Center at Tokyo Tech for DNA sequencing. This work was supported by MEXT Grants-in-Aid for Scientific Research (grant numbers JP26116002, JP18H03984, JP21H04763, and JP20H05925 to HT), JST, the establishment of university fellowships towards the creation of science technology innovation (grant number JPMJFS2112 to H Ito), Uehara Memorial Foundation, Mitsubishi Foundation, and Daiichi Sankyo.

## Author Contributions

H.Ito and K.M. performed experiments; H.Ito K.M. M.U. Y.N. H.Imataka. and H.T. conceived the study, designed experiments, analyzed the results approved the manuscript, and are accountable for all aspects of the work; H.T. supervised the entire project; H.Ito and H.T. wrote the manuscript.

## Supporting Information

**Fig. S1:**
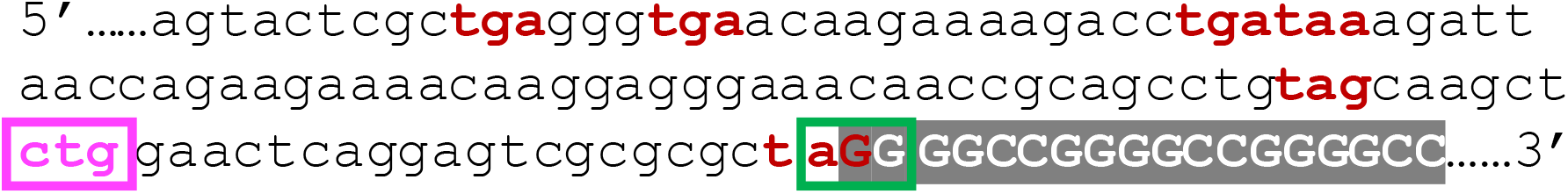
The sequence of the first intron of *C9orf72*. The GGGGCC repeat is represented in a shaded gray box. The CTG codon, positioned at -24 nucleotides (magenta box) from the repeat sequence, was previously identified as the start codon for the GA frame (32–35). The AGG codon, overlapping with the repeat sequence (green box), was previously identified as the start codon for the GR frame (34). Stop codons are present in all frames (red), corresponding to the GP, GP, GR, GR, GA, and GP frames, sequentially arranged from the 5’ to 3’ direction.

**Fig. S2:**
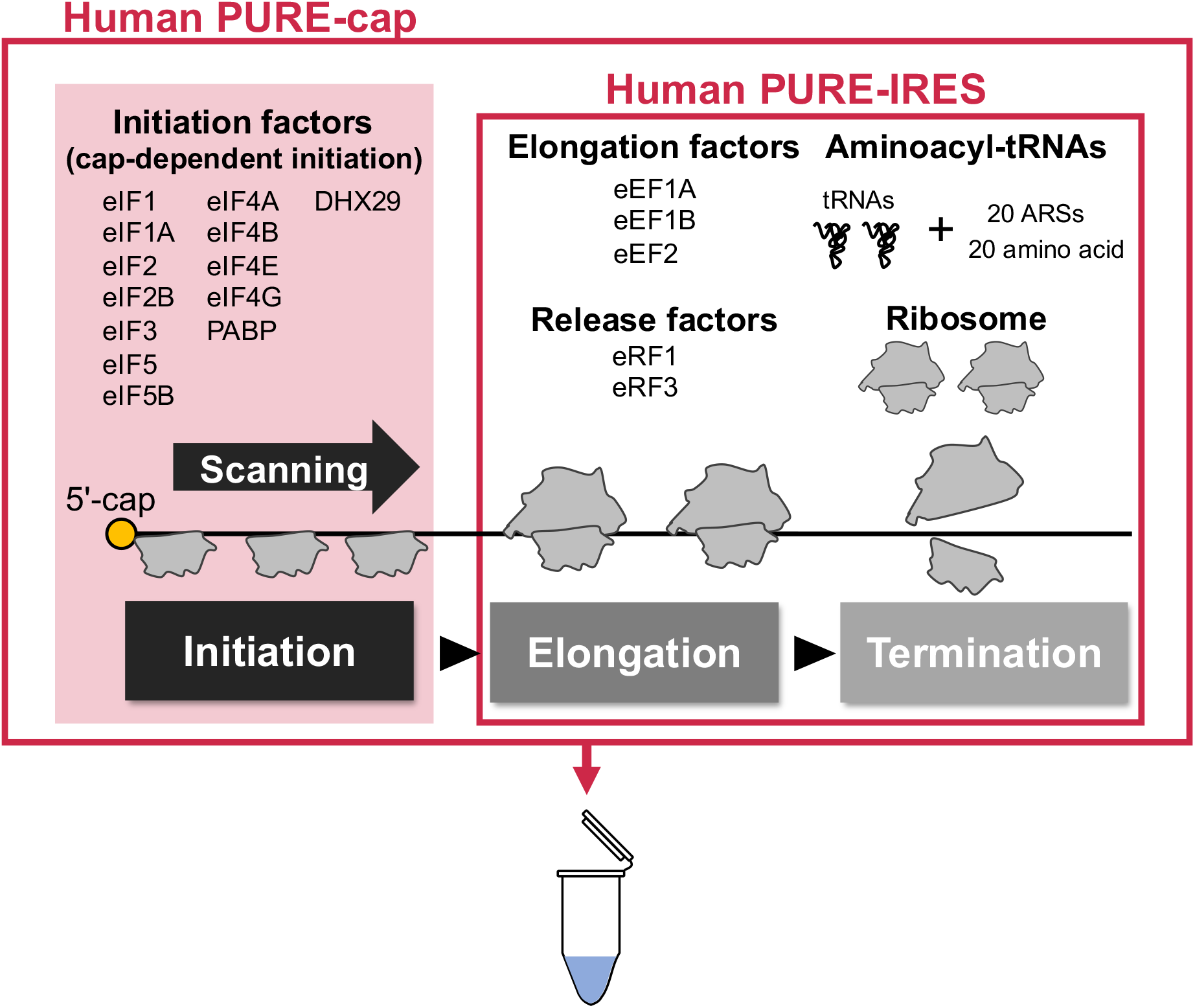
A schematic depiction of the human PURE system. Human PURE-IRES consists of two elementary steps in translation: namely elongation and termination. Human PURE-cap includes eukaryotic initiation factors (eIFs) responsible for scanning-mediated translation initiation, in addition to human PURE-IRES.

**Fig. S3:**
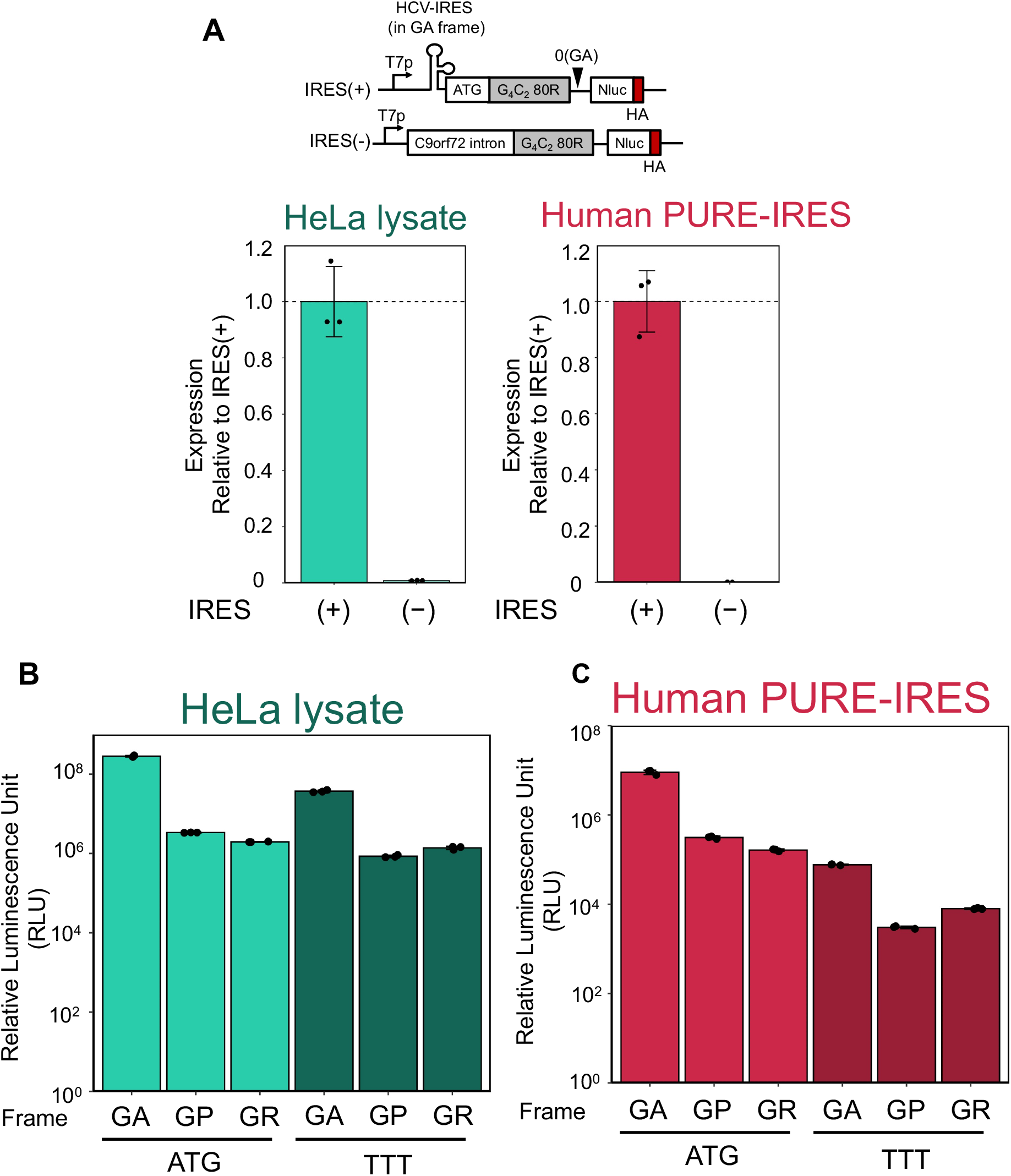
HCV IRES-mediated translation using either HeLa lysate or human PURE-IRES. (*A*) The expression of IRES(-)-(G_4_C_2_)_80_-Nluc and IRES(+) reporters in each in vitro translation system. (*B*, *C*) The absolute values of relative luminescence units (RLU) in HeLa lysate (*B*) or human PURE-IRES (*C*). Error bars indicate standard deviations (±SD) derived from three technical replicates.

**Fig. S4:**
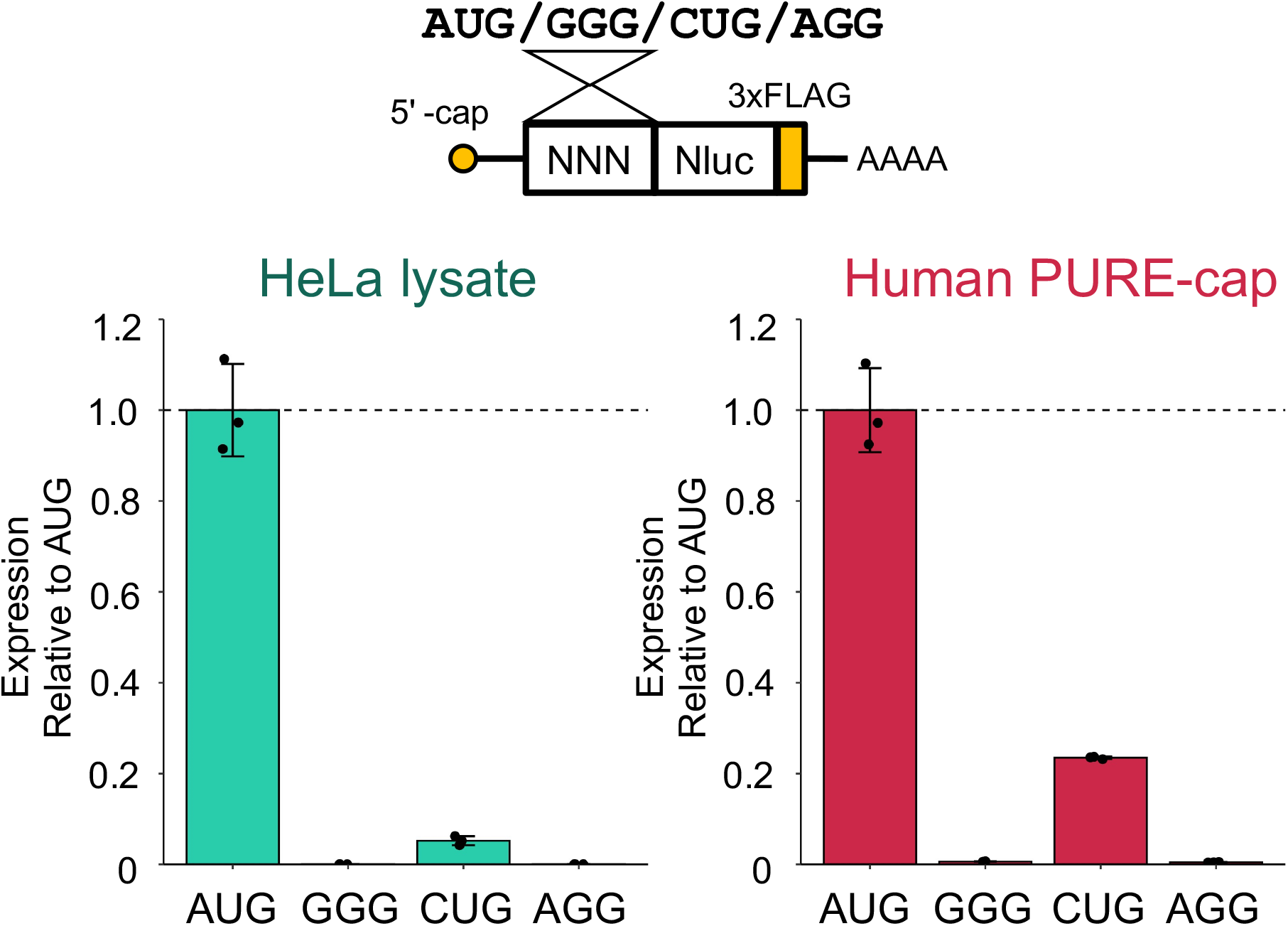
Characteristics of non-AUG translation initiation in either HeLa lysate or human PURE-cap. Relative expression levels of non-AUG translation reporters in each in vitro translation system are shown. The values were normalized with respect to that of AUG-Nluc. Error bars indicate ±SD from three technical replicates.

**Fig. S5:**
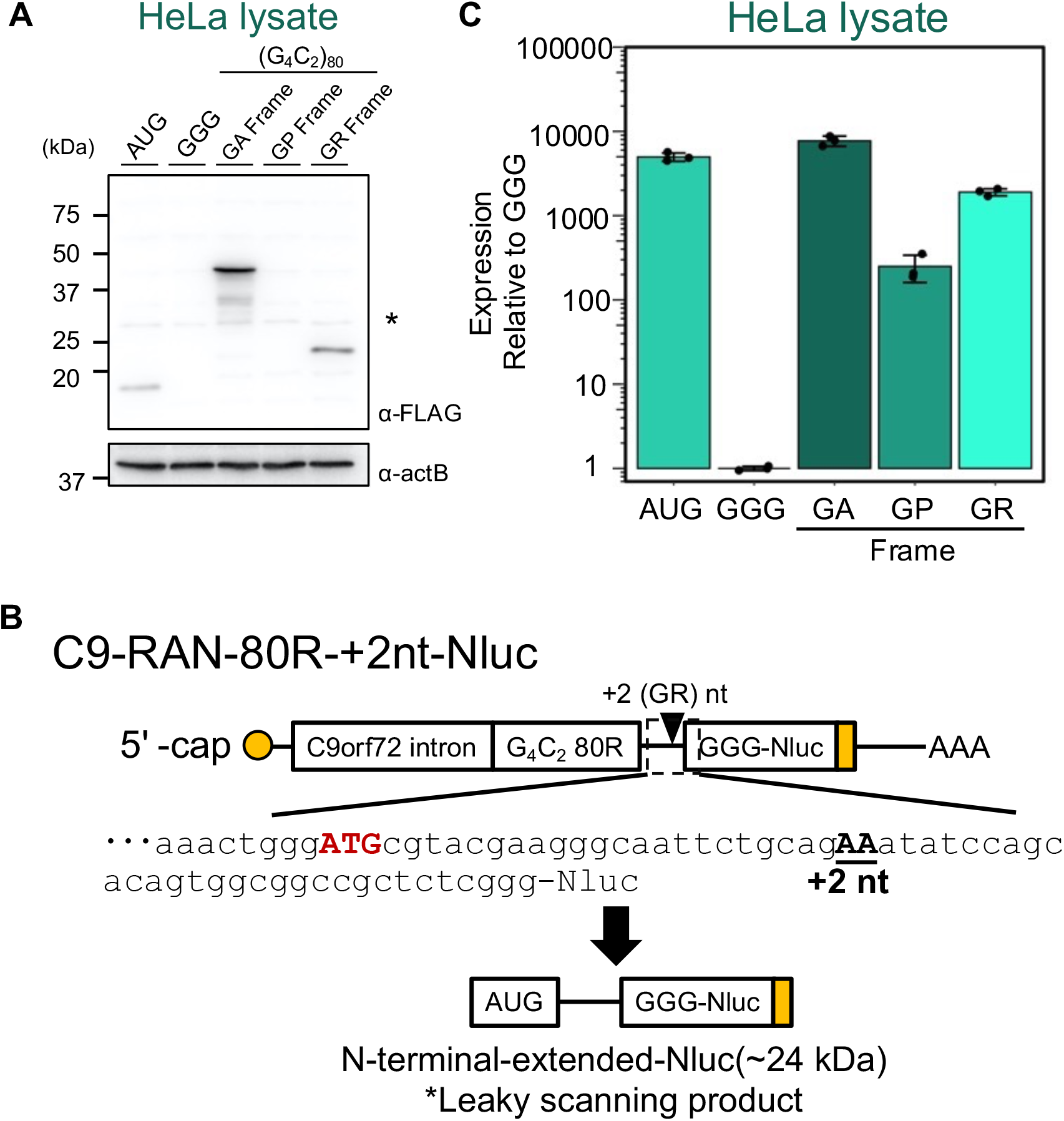
Cap-mediated translation of the C9-RAN reporters in the HeLa lysate. *(A)* An anti-FLAG western blot of the C9-RAN reporter plasmids expressed in the HeLa lysate is shown. An asterisk (*) denotes a non-specific band. *(B)* The nucleotide sequence downstream of the G_4_C_2_ repeat, highlighting an ATG codon potentially associated with a leaky scanning product in the C9-RAN-80R-+2 nt-Nluc reporter. *(C)* Relative expression levels of C9-RAN reporters are presented. The values were normalized to that of GGG-Nluc. Error bars indicate ±SD from three technical replicates.

**Fig. S6:**
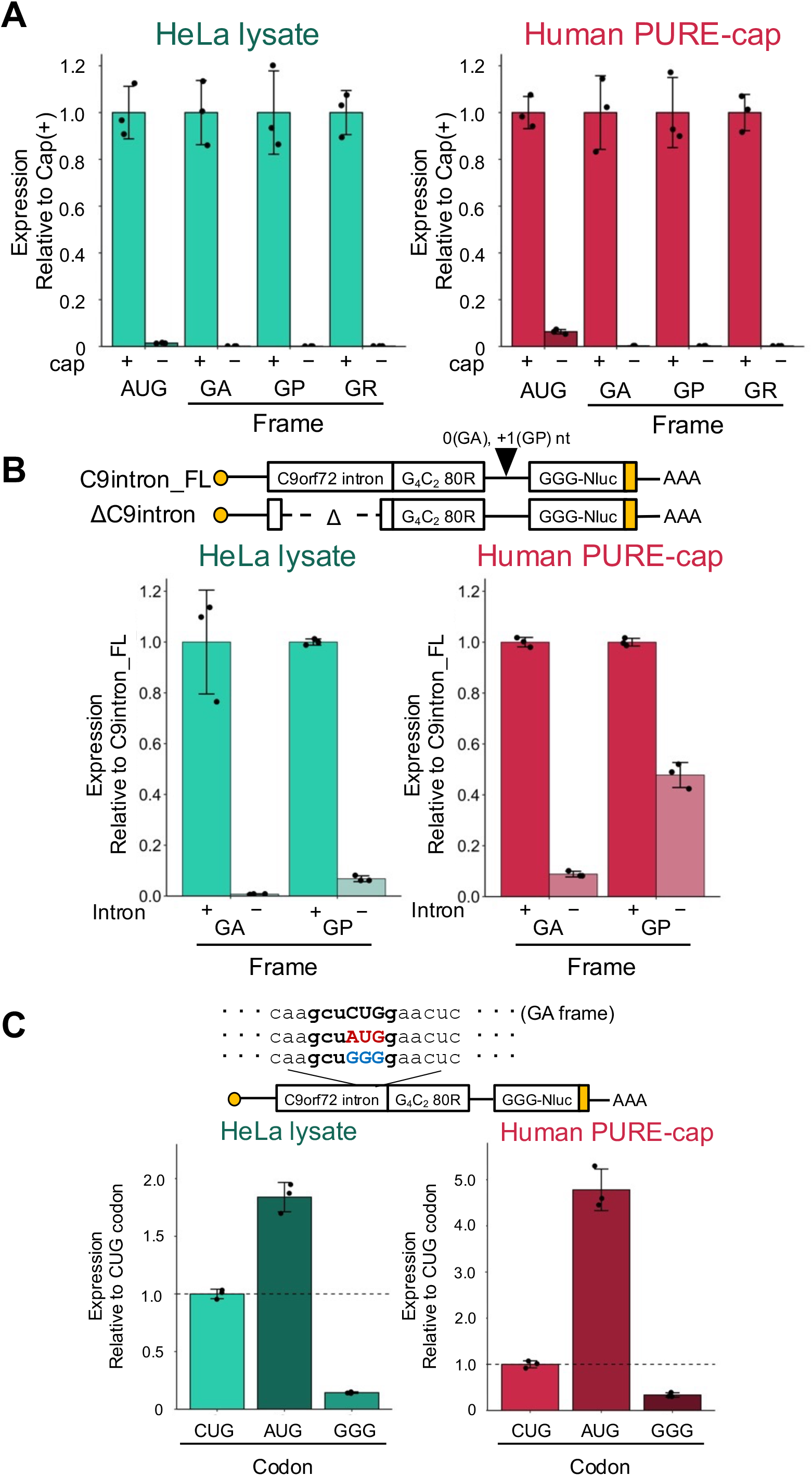
C9-RAN initiates translation from the upstream of the repeat sequence in a 5’-cap-dependent manner. (A) 5’-cap-dependecy: The values were normalized to those of cap(+) C9-RAN reporters. (B) C9 intron dependency: The values were normalized to those of the C9 intron_FL-containing reporters. (C) CUG in C9 intron dependency: The values were normalized to those of the CUG codon in the GA frame located upstream of the G_4_C_2_ repeat. Error bars indicate ±SD from three technical replicates.

**Fig. S7:**
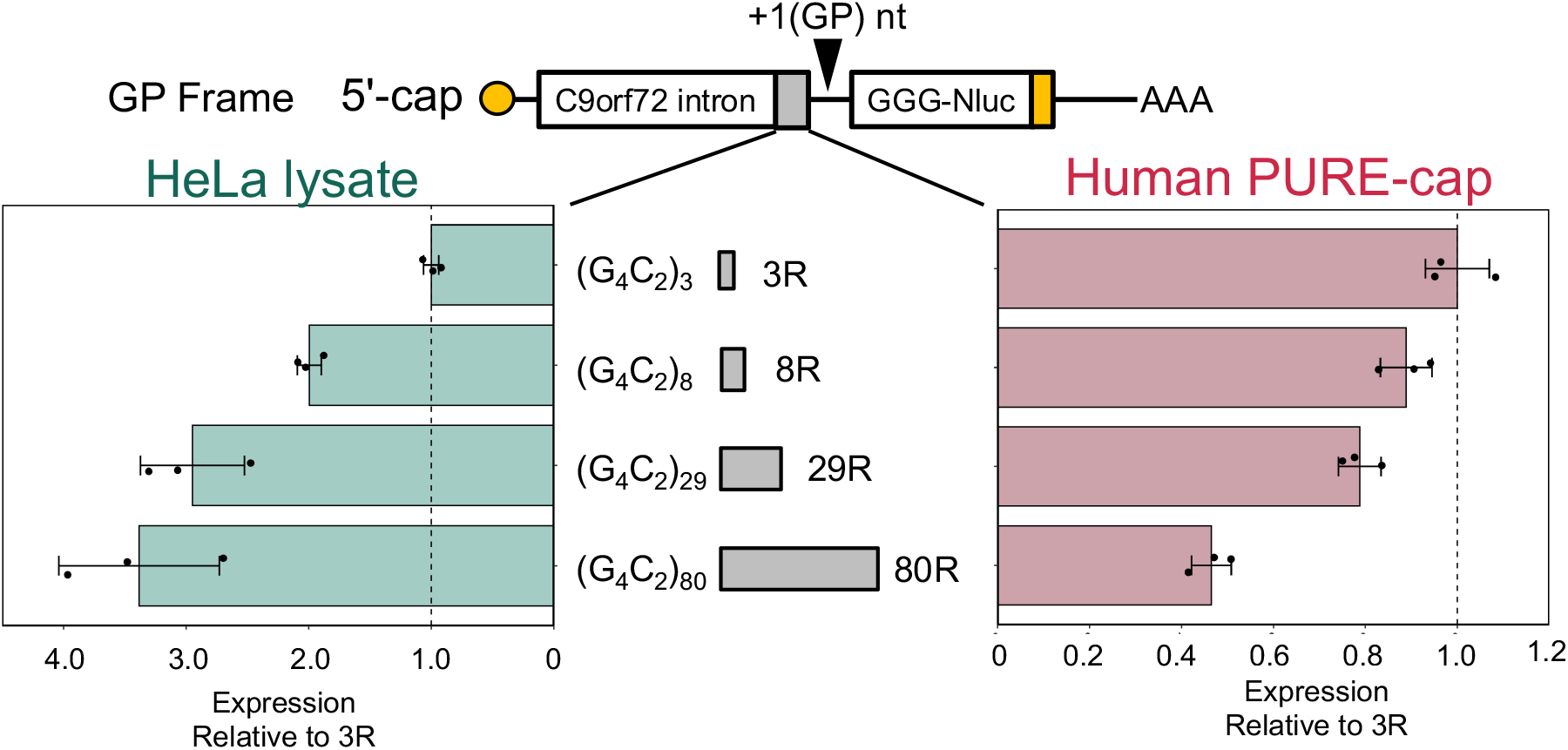
Repeat-length dependent inhibition of C9-RAN (the GP frame) in the human PURE-cap system. Relative expression levels of C9-RAN translation reporters with varying numbers of repeats in each in vitro translation system in the GP frame are presented. The values were normalized to those of 3R-Nluc. Error bars indicate ±SD from three technical replicates.

## References

1. A. J. Hannan, Tandem repeats mediating genetic plasticity in health and disease. Nature Reviews Genetics 2018 19:5 19, 286–298 (2018).

2. I. Malik, C. P. Kelley, E. T. Wang, P. K. Todd, Molecular mechanisms underlying nucleotide repeat expansion disorders. Nature Reviews Molecular Cell Biology 2021 22:9 22, 589–607 (2021).

3. T. Zu, et al., Non-ATG-initiated translation directed by microsatellite expansions. Proc Natl Acad Sci U S A 108, 260–265 (2011).

4. P. E. A. Ash, et al., Unconventional translation of C9ORF72 GGGGCC expansion generates insoluble polypeptides specific to c9FTD/ALS. Neuron 77, 639–46 (2013).

5. T. Zu, et al., RAN Translation Regulated by Muscleblind Proteins in Myotonic Dystrophy Type 2. Neuron 95, 1292–1305.e5 (2017).

6. K. Mori, et al., The C9orf72 GGGGCC repeat is translated into aggregating dipeptide-repeat proteins in FTLD/ALS. Science (1979) 339, 1335–1338 (2013).

7. M. Bañez-Coronel, et al., RAN Translation in Huntington Disease. Neuron 88, 667–677 (2015).

8. E. Soragni, et al., Repeat-Associated Non-ATG (RAN) Translation in Fuchs’ Endothelial Corneal Dystrophy. Invest Ophthalmol Vis Sci 59, 1888–1896 (2018).

9. P. K. Todd, et al., CGG Repeat-Associated Translation Mediates Neurodegeneration in Fragile X Tremor Ataxia Syndrome. Neuron 78, 440–455 (2013).

10. M. DeJesus-Hernandez, et al., Expanded GGGGCC Hexanucleotide Repeat in Noncoding Region of C9ORF72 Causes Chromosome 9p-Linked FTD and ALS. Neuron 72, 245–256 (2011).

11. A. E. Renton, et al., A hexanucleotide repeat expansion in C9ORF72 is the cause of chromosome 9p21-linked ALS-FTD. Neuron 72, 257–268 (2011).

12. T. F. Gendron, et al., Antisense transcripts of the expanded C9ORF72 hexanucleotide repeat form nuclear RNA foci and undergo repeat-associated non-ATG translation in c9FTD/ALS. Acta Neuropathol 3, 829–844 (2013).

13. T. Zu, et al., RAN proteins and RNA foci from antisense transcripts in C9ORF72 ALS and frontotemporal dementia. Proceedings of the National Academy of Sciences 110, E4968–E4977 (2013).

14. A. Jovičič, et al., Modifiers of C9orf72 dipeptide repeat toxicity connect nucleocytoplasmic transport defects to FTD/ALS. Nature Neuroscience 2015 18:9 18, 1226–1229 (2015).

15. Y. J. Zhang, et al., Poly(GR) impairs protein translation and stress granule dynamics in C9orf72-associated frontotemporal dementia and amyotrophic lateral sclerosis. Nat Med 24, 1136–1142 (2018).

16. Y. J. Zhang, et al., C9ORF72 poly(GA) aggregates sequester and impair HR23 and nucleocytoplasmic transport proteins. Nature Neuroscience 2016 19:5 19, 668–677 (2016).

17. S. Mizielinska, et al., C9orf72 repeat expansions cause neurodegeneration in Drosophila through arginine-rich proteins. Science (1979), 1192–1194 (2014).

18. B. Khosravi, et al., Cytoplasmic poly-GA aggregates impair nuclear import of TDP-43 in C9orf72 ALS/FTLD. Hum Mol Genet 26, 790–800 (2017).

19. S. May, et al., C9orf72 FTLD/ALS-associated Gly-Ala dipeptide repeat proteins cause neuronal toxicity and Unc119 sequestration. Acta Neuropathol 128, 485– 503 (2014).

20. K. Mori, et al., Bidirectional transcripts of the expanded C9orf72 hexanucleotide repeat are translated into aggregating dipeptide repeat proteins. Acta Neuropathol 3, 881–893 (2013).

21. A. P. Schuller, R. Green, Roadblocks and resolutions in eukaryotic translation. Nat Rev Mol Cell Biol 19 (2018).

22. N. Sonenberg, A. G. Hinnebusch, Regulation of Translation Initiation in Eukaryotes: Mechanisms and Biological Targets. Cell 136 (2009).

23. R. J. Jackson, C. U. T. Hellen, T. V. Pestova, The mechanism of eukaryotic translation initiation and principles of its regulation. Nat Rev Mol Cell Biol 11 (2010).

24. N. T. Ingolia, L. F. Lareau, J. S. Weissman, Ribosome profiling of mouse embryonic stem cells reveals the complexity and dynamics of mammalian proteomes. Cell 147, 789–802 (2011).

25. 25. N. T. Ingolia, S. Ghaemmaghami, J. R. S. Newman, J. S. Weissman, “Genome-Wide Analysis in Vivo of Translation with Nucleotide Resolution Using Ribosome Profiling” (2009).

26. J. Chen, et al., Pervasive functional translation of noncanonical human open reading frames. Science (1979) 367, 1140–1146 (2020).

27. M. G. Kearse, J. E. Wilusz, Non-AUG translation: A new start for protein synthesis in eukaryotes. Genes Dev 31, 1717–1731 (2017).

28. S. E. Dmitriev, et al., GTP-independent tRNA delivery to the ribosomal P-site by a novel eukaryotic translation factor. Journal of Biological Chemistry 285, 26779–26787 (2010).

29. S. R. Starck, et al., Leucine-tRNA initiates at CUG start codons for protein synthesis and presentation by MHC class I. Science (1979) 336, 1719–1723 (2012).

30. M. A. Skabkin, et al., Activities of Ligatin and MCT-1/DENR in eukaryotic translation initiation and ribosomal recycling. Genes Dev 24, 1787–1801 (2010).

31. Y. Yang, Z. Wang, IRES-mediated cap-independent translation, a path leading to hidden proteome. J Mol Cell Biol 11 (2019).

32. K. M. Green, et al., RAN translation at C9orf72-associated repeat expansions is selectively enhanced by the integrated stress response. Nat Commun 8 (2017).

33. R. Tabet, et al., CUG initiation and frameshifting enable production of dipeptide repeat proteins from ALS/FTD C9ORF72 transcripts. Nat Commun 9, 1–14 (2018).

34. M. Boivin, et al., Reduced autophagy upon C9ORF72 loss synergizes with dipeptide repeat protein toxicity in G4C2 repeat expansion disorders. EMBO J 39, e100574 (2020).

35. Y. Sonobe, et al., Translation of dipeptide repeat proteins from the C9ORF72 expanded repeat is associated with cellular stress. Neurobiol Dis (2018) https://doi.org/10.1016/j.nbd.2018.05.009.

36. W. Cheng, et al., C9ORF72 GGGGCC repeat-associated non-AUG translation is upregulated by stress through eIF2α phosphorylation. Nat Commun 9 (2018).

37. A. Lampasona, S. Almeida, F. B. Gao, Translation of the poly(GR) frame in C9ORF72-ALS/FTD is regulated by cis-elements involved in alternative splicing. Neurobiol Aging 105 (2021).

38. H. M. van ’t Spijker, et al., Ribosome Profiling Reveals Novel Regulation of C9ORF72 GGGGCC Repeat-Containing RNA Translation. RNA (2021) https://doi.org/10.1261/rna.078963.121.

39. K. M. Green, S. L. Miller, I. Malik, P. K. Todd, Non-canonical initiation factors modulate repeat-associated non-AUG translation. Hum Mol Genet 31, 2521– 2534 (2022).

40. Y. Sonobe, et al., A C. elegans model of C9orf72-associated ALS/FTD uncovers a conserved role for eIF2D in RAN translation. Nat Commun 12 (2021).

41. M. Sauert, H. Temmel, I. Moll, Heterogeneity of the translational machinery: Variations on a common theme. Biochimie 114, 39–47 (2015).

42. Y. Shimizu, et al., Cell-free translation reconstituted with purified components. Nature Biotechnology 2001 19:8 19, 751–755 (2001).

43. K. Machida, et al., A translation system reconstituted with human factors proves that processing of encephalomyocarditis virus proteins 2A and 2B occurs in the elongation phase of translation without eukaryotic release factors. Journal of Biological Chemistry 289 (2014).

44. K. Machida, et al., Dynamic interaction of poly(A)-binding protein with the ribosome. Sci Rep 8 (2018).

45. T. Abe, R. Nagai, H. Imataka, N. Takeuchi-Tomita, Reconstitution of yeast translation elongation and termination in vitro utilizing CrPV IRES-containing mRNA. The Journal of Biochemistry 167, 441–450 (2020).

46. A. M. Lancaster, E. Jan, P. Sarnow, Initiation factor-independent translation mediated by the hepatitis C virus internal ribosome entry site. RNA 12 (2006).

47. C. Sellier, et al., Translation of Expanded CGG Repeats into FMRpolyG Is Pathogenic and May Contribute to Fragile X Tremor Ataxia Syndrome. Neuron 93, 331–347 (2017).

48. M. G. Kearse, et al., CGG Repeat-Associated Non-AUG Translation Utilizes a Cap-Dependent Scanning Mechanism of Initiation to Produce Toxic Proteins. Mol Cell 62, 314–322 (2016).

49. T. Zu, Non-ATG-initiated translation directed by microsatellite expansions. Proc. Natl. Acad. Sci. U.S.A. 108, 260–265 (2011).

50. M. G. Kearse, J. E. Wilusz, Non-AUG translation: A new start for protein synthesis in eukaryotes. Genes Dev 31, 1717–1731 (2017).

51. A. Chingakham Ranjit Singh, et al., Human oncoprotein 5MP suppresses general and repeat-associated non-AUG translation via eIF3 by a common mechanism ll Human oncoprotein 5MP suppresses general and repeat-associated non-AUG translation via eIF3 by a common mechanism. Cell Rep 36 (2021).

52. J. D. Cleary, L. P. Ranum, New developments in RAN translation: insights from multiple diseases. Curr Opin Genet Dev 44, 125–134 (2017).

53. J. D. Cleary, A. Pattamatta, L. P. W. Ranum, Repeat-associated non-ATG (RAN) translation. Journal of Biological Chemistry 293 (2018).

54. C.-H. Yu, M.-P. Teulade-Fichou, R. C. L. Olsthoorn, Stimulation of ribosomal frameshifting by RNA G-quadruplex structures. Nucleic Acids Res 42, 1887– 1892 (2014).

55. T. Ishiguro, et al., Regulatory Role of RNA Chaperone TDP-43 for RNA Misfolding and Repeat-Associated Translation in SCA31. Neuron 94, 108–124.e7 (2017).

56. X. Wang, et al., Regulation of HIV-1 Gag-Pol Expression by Shiftless, an Inhibitor of Programmed -1 Ribosomal Frameshifting. Cell 176, 625–635.e14 (2019).

57. M. Kozak, Downstream secondary structure facilitates recognition of initiator codons by eukaryotic ribosomes. Proc Natl Acad Sci U S A 87 (1990).

58. M. G. Kearse, et al., Ribosome queuing enables non-AUG translation to be resistant to multiple protein synthesis inhibitors (2019) https://doi.org/10.1101/gad.324715.119.

59. Y.-J. Tseng, et al., The RNA helicase DHX36/G4R1 modulates C9orf72 GGGGCC hexanucleotide repeat-associated translation. Journal of Biological Chemistry, 100914 (2021).

60. H. Liu, et al., A Helicase Unwinds Hexanucleotide Repeat RNA G-Quadruplexes and Facilitates Repeat-Associated Non-AUG Translation. J. Am. Chem. Soc 143, 7368–7379 (2021).

61. W. Cheng, et al., CRISPR-Cas9 Screens Identify the RNA Helicase DDX3X as a Repressor of C9ORF72 (GGGGCC)n Repeat-Associated Non-AUG Translation. Neuron 104, 885–898.e8 (2019).

62. F. Yuzo, et al., FUS regulates RAN translation through modulating the G-quadruplex structure of GGGGCC repeat RNA in C9orf72-linked ALS/FTD. Elife 12 (2023).

63. S. Mikami, T. Kobayashi, M. Masutani, S. Yokoyama, H. Imataka, A human cell-derived in vitro coupled transcription/translation system optimized for production of recombinant proteins. Protein Expr Purif 62 (2008).

